# How musicality enhances top-down and bottom-up selective attention: Insights from precise separation of simultaneous neural responses

**DOI:** 10.1101/2024.08.23.609277

**Authors:** Cassia Low Manting, Dimitrios Pantazis, John Gabrieli, Daniel Lundqvist

## Abstract

Natural environments typically contain a blend of simultaneous sounds. A significant challenge in neuroscience is identifying specific neural signals corresponding to each sound and analyzing them separately. Combining frequency-tagging and machine learning, we achieved high-precision separation of neural responses to mixed melodies, classifying them by selective attention towards specific melodies. Across two magnetoencephalography datasets, individual musicality and task performance heavily influenced the attentional recruitment of cortical regions, correlating positively with top-down attention in the left parietal cortex, but negatively with bottom-up attention in the right. In prefrontal areas, neural responses indicating higher sustained selective attention reflected better performance and musicality. These results suggest musical training enhances neural mechanisms in the frontoparietal regions, boosting performance via improving top-down attention, reducing bottom-up distractions, and maintaining selective attention over time. Importantly, this work establishes the effectiveness of combining frequency-tagging with machine learning to capture cognitive and behavioral effects with stimulus precision, applicable to other studies involving simultaneous stimuli.

## 2. INTRODUCTION

Analyzing neurophysiological signals generated by multiple concurrent stimuli is challenging due to the difficulty in resolving brain activity to their respective stimuli. In auditory neuroscience, an effective method to achieve this is crucial as simultaneous sounds are commonly encountered in naturalistic environments, such as a cocktail party scenario or an orchestra performance.

In terms of separation precision, few techniques can rival frequency-tagging – a technique which labels neural activity with a unique frequency-based ID tag, allowing it to be identified from a mixture of other activities. Frequency-tagging elicits a neural auditory steady-state response (ASSR) which phase-locks to the envelope of the driving auditory stimulus and is measurable via electrophysiological methods like magnetoencephalography (MEG). Simultaneous neural ASSRs with unique frequencies can thus be reliably isolated and extracted using power spectral density estimation techniques such as Fourier analysis^1,2^. The magnitude and spatial distribution of ASSRs over time offer a powerful, non-invasive means of characterizing distributed neural processing across the brain with exceptional precision.

Due to its unique capability to generate robust and reproducible neural signatures^2,3^, frequency-tagging has found applications in both clinical^4,5^ and experimental research. In cognitive neuroscience, it has been used to investigate intermodal selective attention, demonstrating that cortical ASSRs are enhanced when attention is directed towards an auditory stimulus from a competing visual stimulus^6–8^. Within intramodal auditory attention, ASSR enhancements have also been observed in dichotic paradigms where spatial attention shifted between the left and right ears ^9–11^.

However, in more complex scenarios, frequency-tagging is notoriously difficult to implement, often producing steady-state responses with insufficient signal-to-noise ratio (SNR) needed to capture subtle cognitive or behavioral effects, such as feature-selective auditory attention. This challenge is worsened when multiple sound sources (e.g. voices/instruments) are presented simultaneously due to the suppression of ASSRs that occurs in the presence of simultaneous sounds, yielding inconsistent results^12–14^. An obstacle faced by many ASSR studies seeking higher SNR is that auditory frequency-tagging is extremely intolerant towards small timing inconsistencies (e.g. jitters in stimulus onset time), as out-of-phase signals can easily cancel each other out during averaging. To circumvent this limitation, we operated a specialized auditory setup that achieves near-zero lag in sound delivery (see **Methods**), and recorded brain activity at high temporal resolution using magnetoencephalography (MEG). Moreover, although signal averaging within regions-of-interest is often implemented improve the SNR, even small inter-subject spatial variations can dilute and obscure potential effects when using such univariate approaches. To address this, we developed a unique multivariate decoding method optimized for high-precision ASSR analysis, with enhanced sensitivity to detect subtle cognitive and behavioral modulations.

Auditory training has been shown to induce neuroplastic changes that enhance auditory processing^15–17^. Music training, in particular, enhances higher-order cognitive skills such as pitch perception, auditory attention, and selective listening in noisy environments, with transfer effects to domains like language processing—possibly mediated by overlapping neural networks^17–22^. Prior evidence from our lab demonstrated that the ASSR power correlates positively with individual musicality^23^, suggesting that the ASSR is sensitive to musical training. However, whether this sensitivity extends to its attentional modulation remains unknown. Leveraging the enhanced sensitivity of our multivariate decoding approach, the present study aims to characterize how musicality influences top-down and bottom-up mechanisms across the cortex, contributing to enhanced selective attention.

In this paper, we established the robustness of our signal separation method, with clear evidence from two experiments, showing that our algorithms excelled in their ability to (i) resolve neural signals according to their driving stimuli with exceptional precision, (ii) isolate and extract the effects of top-down and bottom-up selective attention to melody pitch over cortical regions, as well as (iii) achieve this on a continuous scale that correlated significantly with individual musicality and performance scores on selective melody listening tasks.

Overall, results demonstrate that the pattern of brain activity and direction of correlations differ significantly between top-down and bottom-up attention at both sensor and source levels, with compelling evidence highlighting the frontoparietal cortices as key drivers of these differences. The findings also indicate that musical training sustains and enhances selective auditory attention over time through frontal mechanisms, offering valuable insights into how learning and expertise optimize brain function to accomplish cognitive goals.

## 3. RESULTS

### General Task description

Participants listened to two simultaneous melodies (see section 5 for details) of different pitch for a randomized duration of 10 - 15 s per block. The low-pitched melody was frequency-tagged at 39 Hz and the high-pitched melody was frequency-tagged at 43 Hz to elicit corresponding neural responses at 39 and 43 Hz (Fig. 1).

**Fig. 1.**
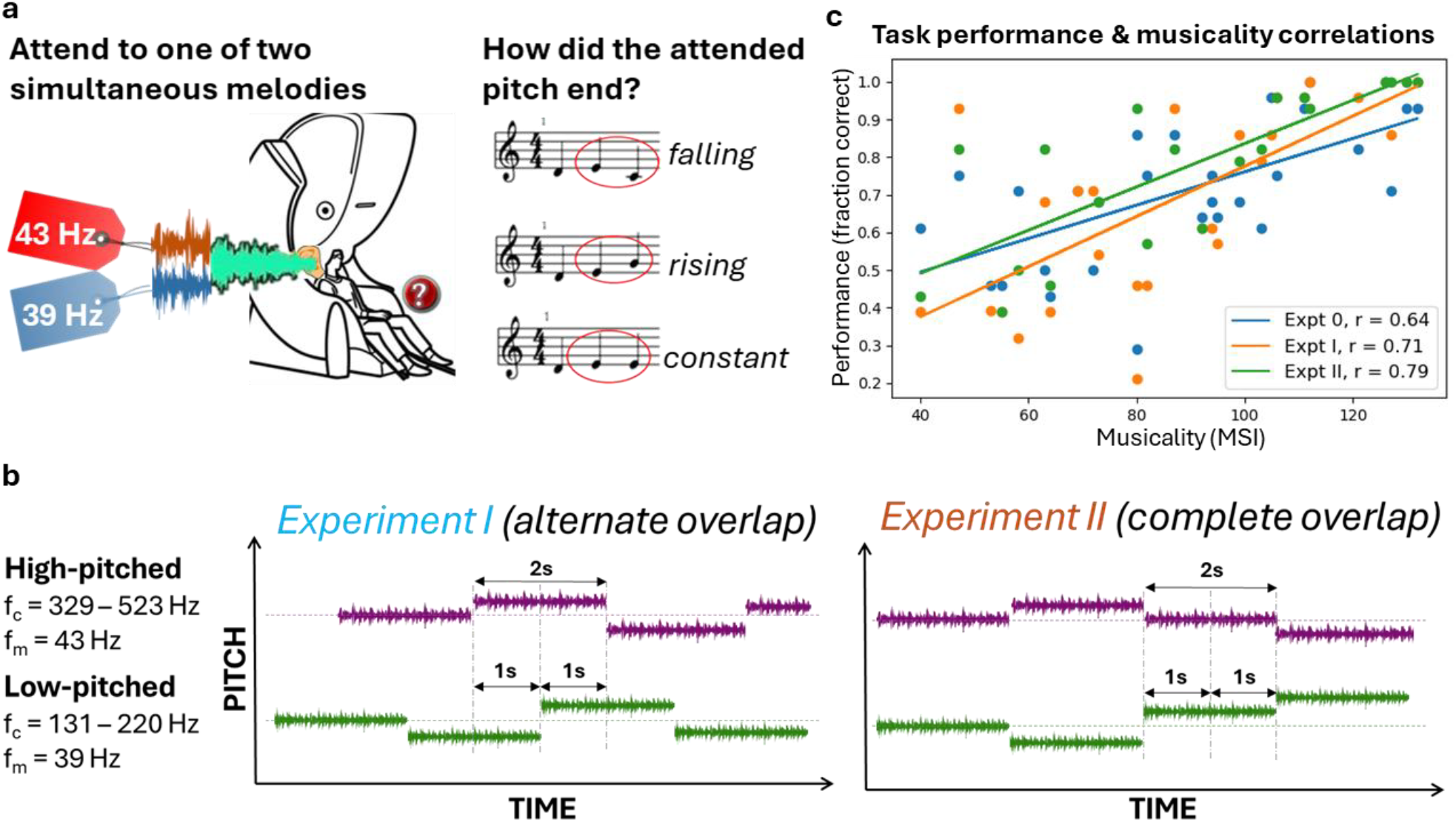
Experimental tasks and behavioral results. **a**, Melody attention task. Participants attended selectively to one of two simultaneous melodies with different pitches. The low-pitched and high-pitched melodies were frequency-tagged at 39 and 43 Hz respectively. When the melody stopped, participants reported the final direction of pitch change for the attended melody, which was either falling, rising or constant. Identical sounds were presented to both ears, ensuring that the melodies could only be distinguished by pitch or timing. **b**, Two variations of the experimental task. Experiment I (left): Alternately overlapping melodies. Tone onsets alternate between the low-pitched and high-pitched melodies. Each new tone induces bottom-up attention towards it, allowing the dissociation of top-down and bottom-up attention effects towards each melody. Experiment II (right): Completely overlapping melodies. Melody tones overlap completely, engaging bottom-up attention simultaneously towards both melodies. All other parameters including modulation frequencies (f_m_) and carrier frequencies (f_c_) were identical to experiment I. **c**, Correlations between participant task performance and musicality. Task performance correlated positively (Pearson correlation, p < 0.001) with musicality across participants in present (Expt I and II) and past experiments (Expt 0). Musicality was measured by the Goldsmiths musical sophistication index^24,25^ (MSI). Notably, the strength of correlation (Pearson r) between musicality and performance increased with task complexity. The least complex task (Expt 0, r = 0.64, N = 28) involved selectively attending to melodies that were completely separated in time and pitch. The moderately complex task (Expt I, r = 0.71, N = 28) required attending to melodies that were partly separated in time and pitch. The most complex task (Expt II, r = 0.79, N = 20) involved selectively attending to melodies that completely overlapped in time and could only be separated by pitch.

For both experiments (experiment I and II), we manipulated top-down selective attention by instructing participants to focus exclusively on either the low-pitched or high-pitched melody. In experiment I, the tone onsets alternate between the low-pitched and high-pitched melodies, drawing an effect of bottom-up attention towards each new tone. Thus, this experimental design allowed us to dissociate top-down and bottom-up attention effects towards each melody. In experiment II, the tone onsets for both melodies occurred concurrently, thereby engaging bottom-up attention simultaneously towards both melodies.

For both experiments, the mean task performance (measured by fraction of correct responses) across participants was above the chance level of 0.33, verifying that selective attention was manipulated accordingly (N_I_=28, m_I_ = 0.70, std_I_=0.25; N_II_=20, m_II_ = 0.77, std_II_=0.21; two-tailed t-tests, p < 0.001;). Furthermore, similar to previous experiments^23,26,27^ (see Supplementary Fig. S1 for description of previous experiments), task performance correlated positively with individual musicality in the current two experiments (Pearson correlation, p < 0.001). Musicality was measured by an adapted version of the Goldsmiths musical sophistication index^24,25^. Notably, the strength of correlation (Pearson r) between musicality and performance increased with task complexity (Fig. 1c). The least complex task (experiment 0, r = 0.64) involved selectively attending to melodies that were completely separated in time and pitch. The moderately complex task (experiment I, r = 0.71) required attending to melodies that were partly separated in time and pitch. The most complex task (experiment II, r = 0.79) involved selectively attending to melodies that completely overlapped in time and could only be separated by pitch.

### Repeated-splitting decoder classified selective attention exclusively at frequency tags

We developed a specialized decoder with heightened sensitivity for ASSRs, utilizing a repeated-splitting classification algorithm (see Fig. 2a and section 5.7 for details). The repeated-splitting classifier was trained on the frequency-tagged neural activity to discriminate between conditions in which attention was directed towards either the low-pitched or high-pitched melody. The area-under-curve (AUC) was computed to measure the classifier’s ability to discriminate between these attention conditions, which was expected to increase with stronger engagement of selective attention.

**Fig. 2.**
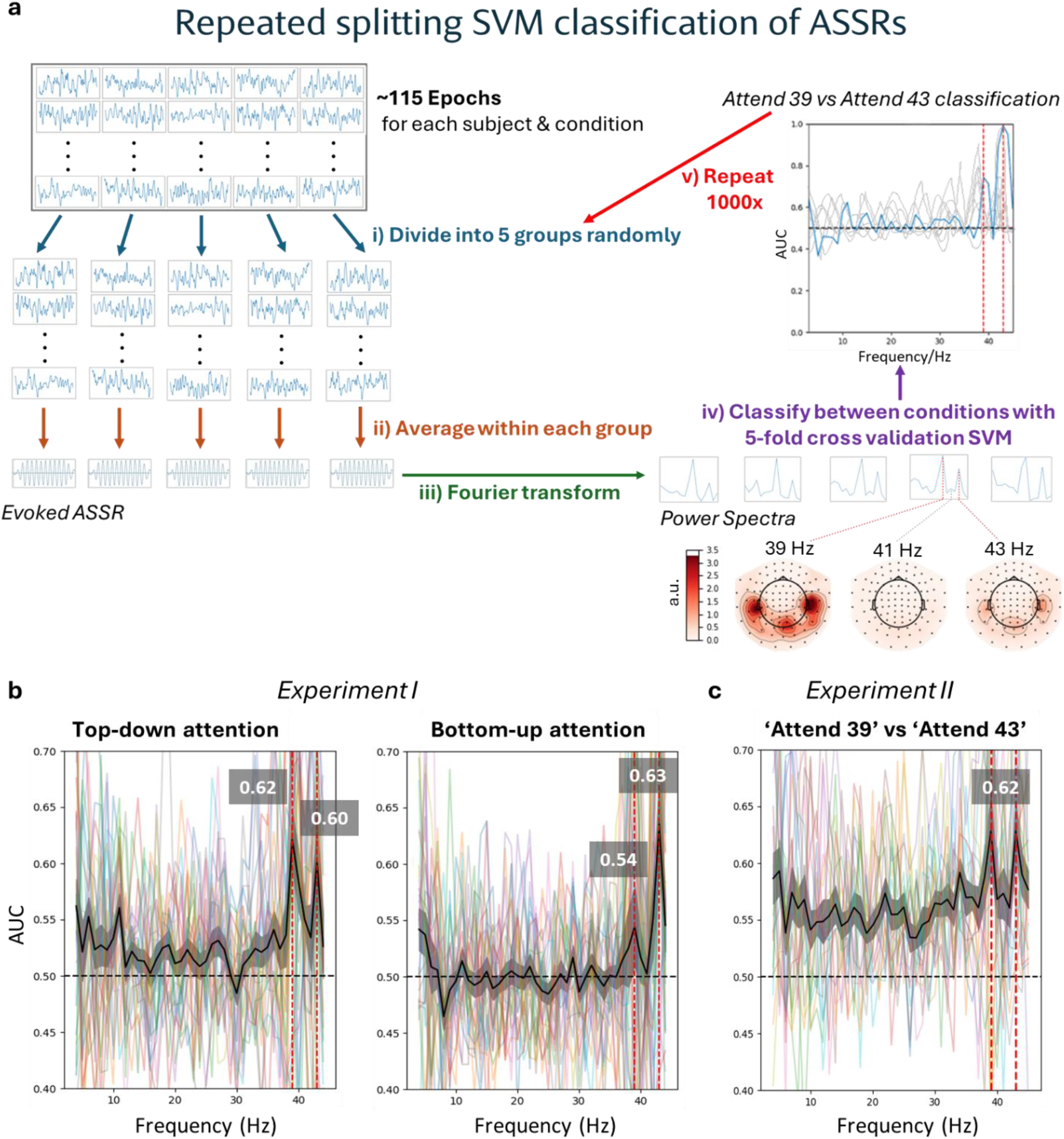
Repeated-splitting classification of selective attention at frequency-tags We trained a classifier on frequency-tagged neural activities to discriminate between conditions where attention was directed towards either the 39 Hz low-pitched or 43 Hz high-pitched melody. **a**, Single-subject repeated-splitting support vector machine pipeline for ASSR classification. For each contrast, epochs per condition were randomly divided into five groups and averaged within each group to produce five evoked ASSRs. Next, the five evoked ASSRs were fourier transformed to acquire five power spectra. For any contrast between two conditions, the corresponding power spectra (5 each condition) were classified via cross-validation, obtaining the area under the receiver operating characteristic curve (AUC) as a measure of discriminability between conditions (chance level = 0.5). The AUC was expected to increase with higher selective attention. Subsequently, we repeated the entire process from the initial group division step 1000 times and computed the mean AUC across repetitions. Note that the AUC was computed independently for each frequency from 4 - 45 Hz. For source analysis, the evoked ASSRs were localized to the cortical surface before fourier transformation (step iii). For each region of interest (ROI), only sources within the ROI were included for classification, combining the power at 39 and 43 Hz in feature space. As in sensor analysis, the mean AUC across 1000 repetitions was computed. Below the power spectra, MEG gradiometer topographies of a single fold input to the classifier demonstrated dominant auditory cortical power precisely tagged to 39 and 43 Hz, but not at the adjacent 41 Hz. For visualization, the power at each frequency was normalized to the mean power across all frequencies and the subject grand average for a single condition in Experiment I is shown. Units are arbitrary (a.u.). **b and c**, Peak AUC values were specifically observed at the stimulus frequency tags of 39 and 43 Hz (red vertical lines), but not at other frequencies. For each plot, the mean across participants is shown in black, with the AUC peak values at 39 and 43 Hz displayed in gray boxes above. **b**, Experiment I (N = 25). Selective attention peaks were observed at 39 and 43 Hz for both top-down (left) and bottom-up attention (right). **c**, Experiment II (N = 20). Similar selective attention peaks were observed at 39 and 43 Hz. These results confirm that we have successfully extracted neural responses at their predefined frequency-tags and captured their attentional modulation. Shading indicates standard error of the mean (s.e.m.).

For experiment I, peak discrimination of the classifier was observed specifically at the set stimulus frequency tags of 39 and 43 Hz, but not at other frequencies. This applied to both top-down (AUC = 0.62 (39 Hz), 0.60 (43 Hz); Fig. 2b – Left) and bottom-up attention (AUC = 0.54 (39 Hz), 0.63 (43 Hz); Fig. 2b – Right). Similarly for experiment II, peak discrimination was observed at 39 and 43 Hz (AUC = 0.62 (39 & 43 Hz), obtained from the higher AUC between the early half and late half of the tone at each frequency; Fig. 2c). The individual tone halves, however, did not exhibit any discrimination peak at the modulation frequencies (Supplementary Fig. S2). These results confirm that we can reliably extract stimulus-specific neural responses at their predefined frequency-tags (i.e. 39 & 43 Hz), and that these responses are sufficiently sensitive to capture the effects of selective attention.

### Bottom-up attention triggers temporal processing; Top-down attention recruits frontal mechanisms

We proceeded to examine how different brain regions are recruited in selective attentional processes, focusing on cortical areas that have been observed to contain ASSR sources in our previous studies^23,26,27^. Six regions of interest, namely the orbital gyrus (OrG), superior temporal gyrus (STG), and inferior parietal lobe (IPL) at each hemisphere, were identified and demarcated with a predefined atlas (Brainnetome Atlas^28^).

Among these six regions, bottom-up attention was discriminated above the chance level of 0.5 only at the right STG, while top-down attention was discriminated above chance at all regions (p < 0.01 for all significant cases, n = 10000, one-tailed permutation test, false discovery rate (FDR)-corrected for 12 tests) except the right IPL (Fig. 3). At the left and right OrG, the classifier could discriminate top-down attention significantly better than bottom-up attention (p_left_ < 0.01, p_right_ < 0.001, n = 10000, two-tailed permutation test, FDR-corrected for 6 tests). These results are in agreement with current literature^29–32^ asserting that bottom-up attention is triggered by lower-level automatic sensory mechanisms situated predominantly in the primary cortices such as the STG, while top-down attention recruits higher-level executive mechanisms located in the prefrontal cortex (Cohen 2014; Foster et al. 1994; S. Huang et al. 2012; Plakke and Romanski 2014).

**Fig. 3.**
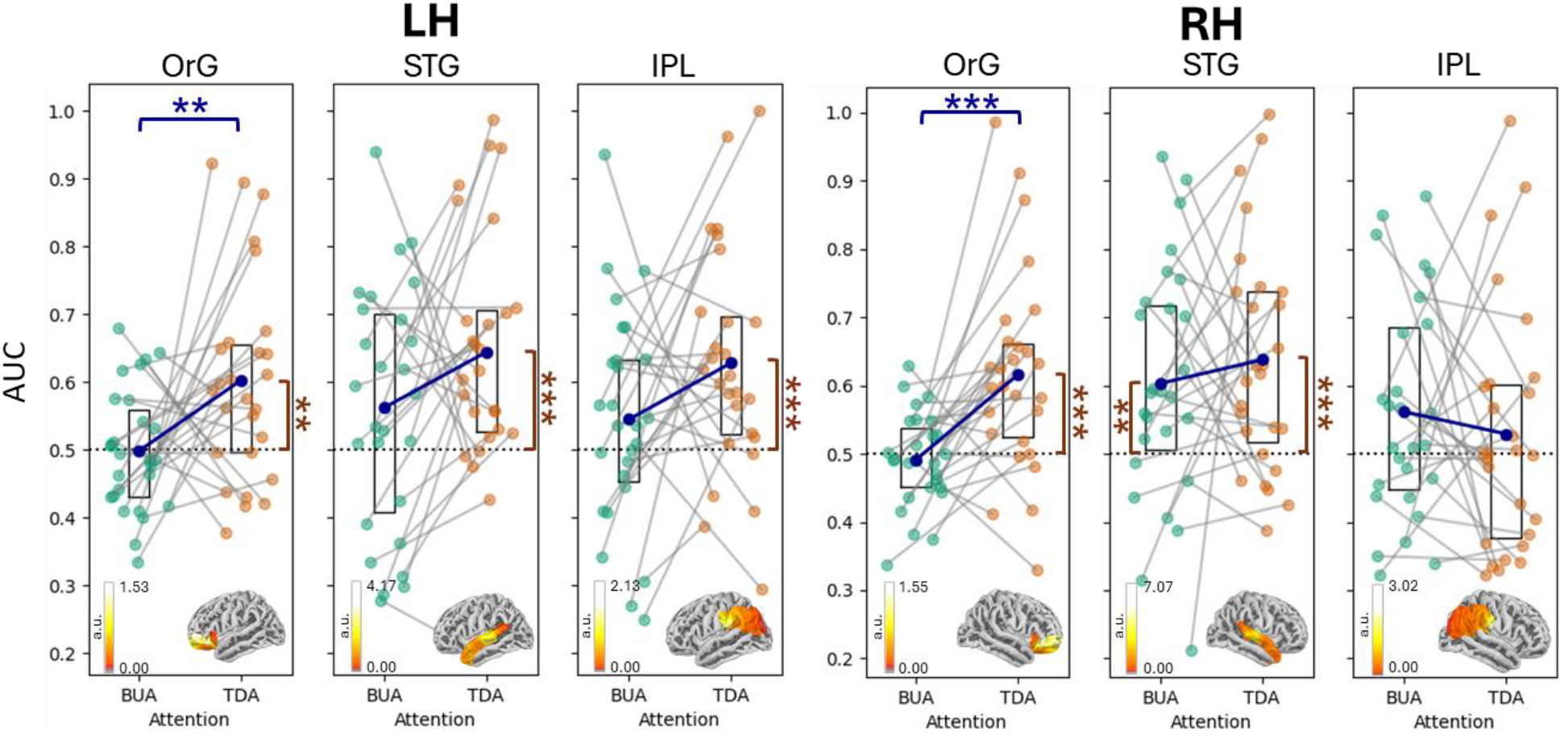
Top-down and bottom-up selective attention across cortical regions Across participants (individual markers; N = 27), bottom-up attention was significantly discriminated above chance (dotted line) only at the right superior temporal gyrus (STG), while top-down attention was discriminated above chance at all regions except the right inferior parietal lobe (IPL). Significance levels above chance are marked by black asterisks beside the corresponding attention condition on the x-axis. The y-axis denotes the AUC and is identical for all subplots. Box plots outline the 25th – 75th percentiles of the data, with the center dot indicating the mean. At the left and right orbital gyri (OrG), top-down attention was discriminated significantly better than bottom-up attention (blue brackets and asterisks). These results support the notion that bottom-up attention is triggered by lower-level automatic sensory mechanisms situated predominantly in the primary cortices such as the STG, while top-down attention recruits higher-level executive mechanisms located in the prefrontal cortex. All p-values are computed with permutation tests and false discovery rate (FDR)-corrected. ***p<0.001, **p<0.01, *p<0.05. Neural activity illustrating cortical power within each region-of-interest is displayed over the standard brain at the bottom of each subplot. For visualization, the subject grand average activity at 39 Hz was normalized to the mean power across all frequencies for a single condition. Units are arbitrary (a.u.).

### Top-down and bottom-up attention show opposing correlations with musicality and performance

To examine the relationship between selective attention, musicality, and task performance, we correlated the AUC with participants’ musicality scores as well as task performance. For musicality, a positive correlation with top-down attention was observed already at sensor level in experiment I (Pearson correlation, r = 0.42, p < 0.05; Supplementary Fig. S3a).

We proceeded to investigated which ASSR sources underlie this correlation, with our primary analysis centered on sources in the IPL, a region that has shown correlations with musicality in our previous study^23^. Indeed, the left IPL exhibited a significant positive correlation between top-down attention and task performance (Pearson correlation, r = 0.49, p < 0.01; Fig. 4a - Left). Further analysis of the remaining regions revealed that this correlation effect was also driven by the left OrG (Pearson correlation, r = 0.40, p < 0.05) and right STG (Pearson correlation, r = 0.39, p < 0.05), albeit to a smaller extent (Supplementary Fig. S3c).

**Fig. 4.**
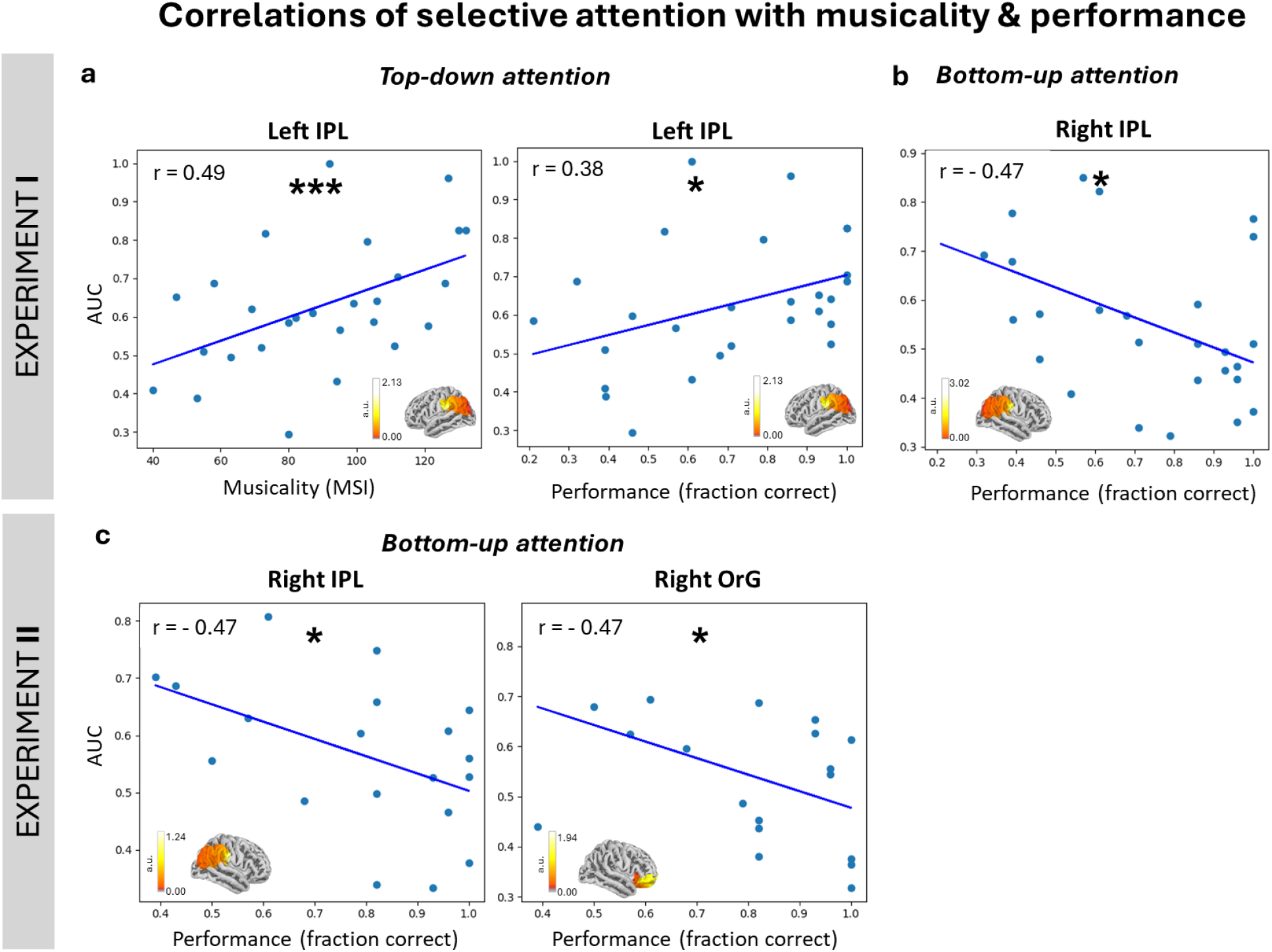
Correlations of selective attention with musicality & performance. Scatterplots showing correlations between classifier area-under-curve (AUC) values against individual musicality or task performance. The AUC is plotted in the y-axis for all subfigures and reflects the degree of selective attention. Neural activity illustrating cortical power within each region-of-interest is displayed over the standard brain at the bottom of each subplot. For visualization, the subject grand average activity at 39 Hz was normalized to the mean power across all frequencies for a single condition. Units are arbitrary (a.u.). **a**, In experiment I, top-down attention at the left inferior parietal lobe (IPL) shows positive correlations with both musicality (left; Pearson correlation, r=0.49, p<0.01) and performance (right; Pearson correlation, r=0.38, p<0.05) across 27 participants. **b**, Across the same participants, bottom-up attention correlated negatively with performance at the right IPL (Pearson correlation, r = -0.47, p < 0.05). **c**, Similarly, in experiment II, bottom-up attention correlated negatively with performance at the right IPL and right orbital gyrus (OrG) across 19 participants (Pearson correlation, r = -0.47, p < 0.05 for both regions).

For task performance, a positive correlation with top-down attention was also observed at the left IPL in experiment I (Pearson correlation, r = 0.38, p < 0.05; Fig. 4a - Right). Contrastingly, the correlation between bottom-up attention and task performance was negative at the right IPL (Pearson correlation, r = -0.47, p < 0.05; Fig. 4b). This negative correlation, although weaker, was already observed at sensor level (Pearson correlation, r = -0.45, p < 0.05; Supplementary Fig. S3b). Furthermore, in experiment II, the early half of the tone showed a negative correlation between bottom-up attention and task performance, but not the late half. This effect was also observed at the right IPL (Pearson correlation, r = -0.47, p< 0.05; Fig. 4c - Left), like in experiment I, as well as at the right OrG (Pearson correlation, r = -0.47, p < 0.05; Fig. 4c - Right). The early half of the tone was thought to elicit a stronger bottom-up attention effect (on both melodies) due to the perceptually salient change in pitch. In contrast, the late half, being a continuation of the same tone, would not strongly draw bottom-up attention. Hence, we infer that the negative correlation manifested from the effect of bottom-up attention rather than top-down attention which was unlikely to differ systematically between the two halves of the tone. Success in performing the experimental task requires participants to actively direct top-down selective attention towards the attended melody, whilst inhibiting bottom-up attentional diversions towards task-irrelevant, salient pitch changes in the competing melody. Consequently, successful inhibition suppresses the effect of bottom-up attention on neural activity, leading to poorer classification. Taken together, the results from both experiments provide compelling evidence that enhancements in selective attention across individuals, likely from musical training, occur by enhancing top-down mechanisms while suppressing bottom-up mechanisms in the frontoparietal regions.

### Musical training sustains selective attention mechanisms in the prefrontal cortex, enhancing performance

To investigate temporal dynamics of selective attention, we employed a sliding time window to calculate AUC values over the 2s tone duration in experiment II. The time of peak selective attention, corresponding to maximum AUC, appeared to follow a bimodal distribution across participants, with a separation at around 0.5 s (Fig. 5a). Participants were thus categorized into two equal groups based on their time of peak selective attention: Early attendees, whose selective attention peaked before 0.5 s from tone onset, and late attendees, whose selective attention peaked after 0.5 seconds. We found that late attendees performed significantly better at the task than early attendees (Δ = 0.19, p < 0.05, n = 10000, two-tailed permutation test; Fig. 5b – Right). Moreover, late attendees tended to be more musical than early attendees, even though the difference nearly missed significance (Δ = 34.3, p = 0.057, n = 10000, two-tailed permutation test; Fig.6b – Left).

**Fig. 5.**
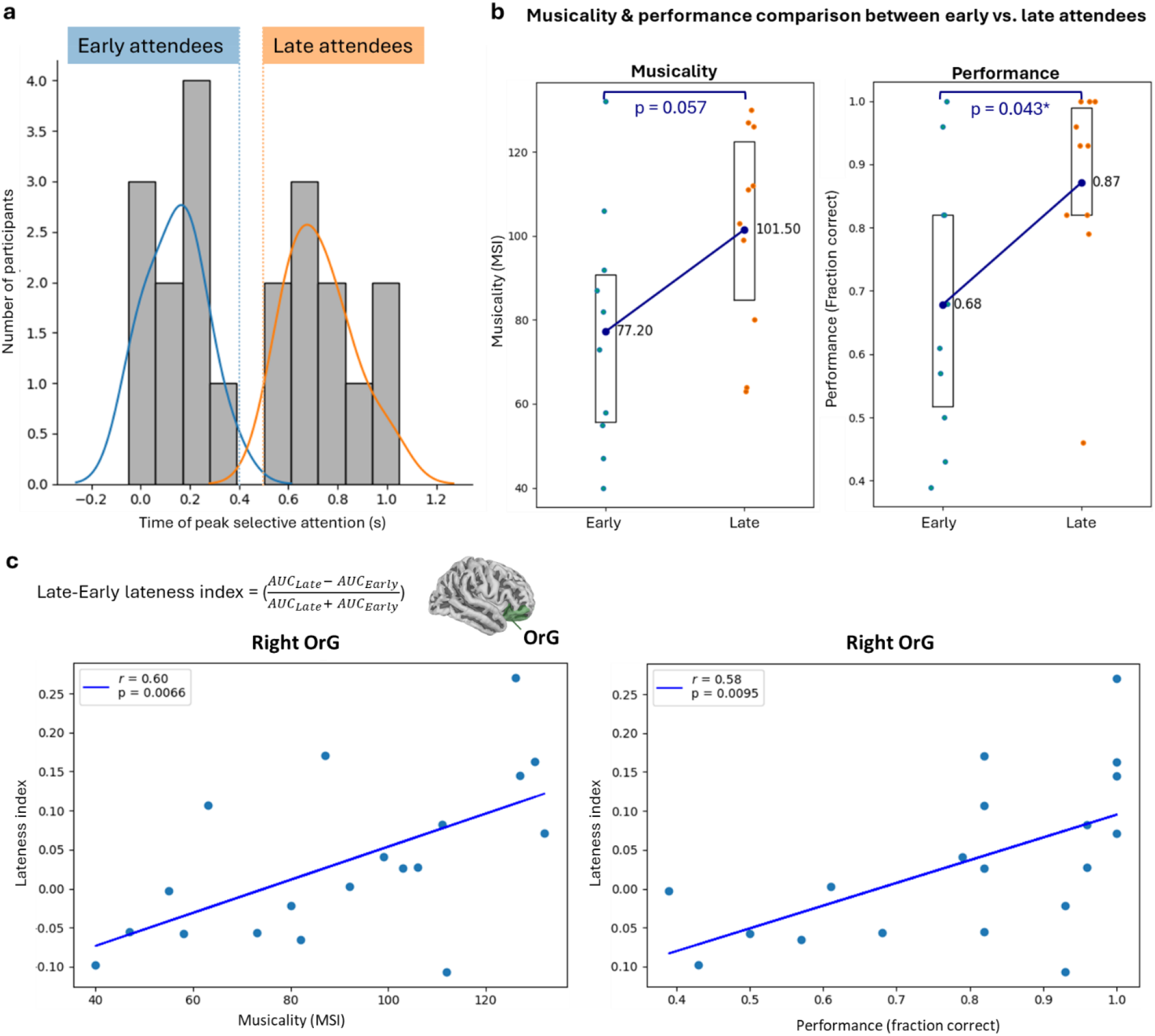
Musicality and performance across early and late attendees. **a**, Histogram of the time of peak selective attention across participants (N = 20). We computed the classifier AUC over the 2 s tone duration using a sliding time window and extracted the time of peak selective attention, corresponding to maximum AUC, for each participant in experiment II. The distribution appeared to be bimodal, dividing participants equally into “early” (blue) or “late” (orange) attendees depending on whether their attention peaked before of after 0.5 s respectively. Colored lines represent normal distributions fitted to each group of attendees. **b**, Musicality (left) & performance (right) comparison between early vs. late attendees. Late attendees performed significantly better (Δ = 0.19, p < 0.05, n = 10000, two-tailed permutation test) at the task than early attendees. Moreover, late attendees tended to be more musical than early attendees, even though the difference nearly missed significance (Δ = 34.3, p = 0.057, n = 10000, two-tailed permutation test). Box plots represent the 25th – 75th percentiles of the data, with the center dot indicating the mean. **c**, Correlational analysis of lateness index with musicality (left) and performance (right). For each participant, we computed a ‘lateness index’ that reflects the relative strength of selective attention in the late half of the tone compared to the early half. The lateness index correlated positively with both musicality and task performance at the right orbital gyrus (OrG, colored green over the adjacent standard brain) (Pearson correlation, p < 0.01 for musicality & performance). Taken together, these results suggest that musical training sharpens the neural mechanisms for sustaining or improving auditory selective attention over time, particularly in the right prefrontal cortex, thereby boosting task performance. The locations of the OrG, STG, and IPL are indicated in Fig. 4d.

At source level, we compared selective attention in the early half of the tone to the late half. To quantify the difference, we computed a ‘lateness index’ for each participant, reflecting the relative strength of selective attention in the late half compared to the early half (Fig. 5c). Our findings at the right OrG revealed positive correlations between the lateness index and both musicality as well as task performance across participants (Pearson correlation, p < 0.01 for musicality & performance; Fig. 5c). Essentially, participants who had better musical skills and task performance paid higher selective attention during the late half compared to the early half of the tone. Conversely, participants who were less musical and performed poorly could not maintain selective attention throughout the tone duration. These results suggest that musical training sharpens the neural mechanisms for sustaining or improving auditory selective attention over time, particularly in the right prefrontal cortex, thereby boosting task performance.

## 4. DISCUSSION

The separation of simultaneous neural responses presents a significant challenge to experimental neuroscience, especially when disentangling subtle cognitive and behavioral effects from complex auditory scenes. By combining the unparalleled separational precision of frequency-tagging with a repeated-splitting classification approach, we isolated the effect of selective attention to a target melody within a mixture of two simultaneous melodies, further demonstrating the implications of individual task performance and musicality, both at sensor and source levels. The combined findings across two experiments demonstrate that our implementation of frequency-tagging in simultaneous stimuli is both reliable and robust. These results elucidated the differential recruitment of cortical regions in bottom-up and top-down attentional processes, aligning with established theories of attentional control which posit that top-down attention engages higher-level, goal-directed executive mechanisms orchestrated by prefrontal regions, while bottom-up attention is driven by automatic lower-level sensory mechanisms predominantly situated in the primary cortices^29–32^. To the best of our knowledge, this is the first successful classification of simultaneous ASSRs based on varying cognitive states and behavior. This breakthrough opens up new possibilities for highly precise separation of simultaneous neural responses with frequency-tagging to study cognitive and behavioral effects.

Frequency-tagging enables the identification and separation of mixed brain responses stemming from simultaneous sounds, which are ubiquitous in our everyday environment. This facilitates the use of research environments that more accurately reflect the complexity of natural soundscapes, ensuring that experimental findings are more applicable to real-life cognitive phenomena. Our approach of applying frequency-tagging at full modulation depth, while eliciting stronger signals, noticeably deteriorates sound quality and naturalness. Decreasing the modulation depth mitigates the degradation in sound quality but may render the ASSRs too weak for extracting cognitive effects within realistic time constraints^1,2^. Therefore, it is imperative for future frequency-tagging endeavors to work on finding an optimal balance between minimizing modulation depth and preserving sufficient signal strength for capturing desired effects. A potential limitation of this work is that replicating our auditory setup (see *Stimulus presentation* under **Methods**) to achieve near zero-lag is tedious and costly, requiring the purchase and installation of new specialized hardware and software if not already accessible. As a more implementable alternative, we recommend recording the auditory output into a separate channel (MISC) of the MEG system to precisely track the time of stimulus delivery. Imperatively, researchers should take into account the Nyquist frequency requirements to capture the auditory signal with sufficiently high fidelity by setting the sampling rate to at least twice the maximum carrier frequency of the auditory stimulus.

Our study provides compelling evidence that top-down and bottom-up attentional mechanisms are differentially influenced by musicality as well as task performance. We found a positive correlation between top-down attention and both musicality and task performance, particularly in the left IPL. This suggests that musical training enhances top-down attentional mechanisms which are crucial for goal-directed behavior and cognitive control, leading to better task performance. Conversely, bottom-up attention showed a negative correlation with task performance in the right IPL, indicating that higher sensitivity to distractions via bottom-up attentional mechanisms may impede performance. Together, these insights contribute to a more nuanced understanding of the neural substrates underlying selective attention, highlighting the distinct yet complementary roles of the left and right parietal cortices in top-down and bottom-up attention. Consistent with our hypothesis, the involvement of the parietal cortex is unsurprising as previous studies from our lab and others^23,33–35^ have demonstrated its role in musical training and the ability to perform music-related tasks.

Similar to the IPL, correlations between selective attention and task performance were also observed in the right OrG, but only under the more demanding condition where the melodies fully overlapped (Experiment II), and not in the partially overlapping condition (Experiment I). Task difficulty modulates the engagement of higher cognitive functions such as selective attention and pitch discrimination, altering activity within frontoparietal neural networks^36–38^. Thus, we postulate that the correlation effects in the OrG emerged only when task demands were sufficient to strongly engage the neural mechanisms involved, thereby exposing individual differences in aptitude. Further evidence for the right OrG’s role in musicality and selective attention ability was revealed through the analysis of temporal attention dynamics. Participants whose selective attention peaked later during the tone performed significantly better and were more musical than those who peaked earlier. Subsequent correlational analysis corroborated with these results, revealing that the stronger attention was during the late tone half, relative to the early tone half, the more musical and better performing the participant was. Our results corroborate longstanding literature establishing the prefrontal cortex as a center for attentional control^29,31,32^, aligning with the hypothesis that musical training sharpens neural mechanisms for sustaining or improving selective attention over time, thereby boosting task performance.

By tracking how the modulation of attention towards each melody varies over time with our sliding window analysis, we can infer the attentional strategies employed by individual participants. Although beyond the scope of this paper, it is fascinating that we found correlations between the attentional modulation of the 39 and 43 Hz melodies overtime, particularly among participants with higher musicality (Supplementary Fig. S4). The data also suggests that these participants tended to perform better, although the difference was not significant. These observations might indicate that musical participants employ an organized strategy to manage attention effectively towards simultaneous sounds, contributing to their better performance. Using a simple index like the Kendall’s Tau, significant correlations between the 39 Hz and 43 Hz attentional modulation curves only emerged in six participants (out of 20). Visual inspection of the individual participant curves revealed more complex correlations with time lags between the 39 and 43 Hz curves for many other participants (Supplementary Fig. S5). Although outside our primary expertise, we believe that it is possible to characterize these complex correlations given a more appropriate index.

Overall, this study contributes to a deeper understanding of the neural substrates underlying selective attention and highlights the potential of musical training as a tool for cognitive enhancement. Further research could expand on these findings by exploring whether instrument-specific musical training impedes attentional control away from the trained instrument tones (e.g., whether a violinist finds it more challenging to ignore a violin tone), and elucidating the mechanisms involved. Additionally, biochemical techniques such as magnetic resonance spectroscopy can provide insights into the neurochemical basis, possibly involving GABA and glutamate neurotransmitters^39–43^, that differentiate selective attention control between high and low performers, or musicians and non-musicians. Finally, while most studies in the field focus mainly on the primary auditory cortex^13,44,7,45,6,9^, where the strongest ASSR sources are situated, our current and past works^23,26,27^ have demonstrated that secondary sources in the frontoparietal regions can better capture cognitive effects such as selective attention and their correlation with behavioral measures (e.g. musicality, task performance). Given their relevance and usefulness, we strongly encourage future research to include these secondary ASSR sources in their analysis. Although the focus of our current work is on auditory attention, frequency-tagging has been used in a diverse range of other disciplines, such as working memory^46–48^, language^49^, aging^50^, cognitive development^51^ and impairment^52^, pain^53–55^, Alzheimer’s disease^52^, schizophrenia^56^ and bipolar disorder^57,58^, as well as sensory modalities including tactile^59,53,54,60^ and vision^61,46,62,48^. Importantly, our repeated-splitting machine learning algorithms can be readily applied to these different fields, enabling researchers to leverage frequency-tagging’s unique ability to separate and precisely trace simultaneous neural signals back to their stimuli, at the same time extracting cognitive and behavioral effects with high-sensitivity.

## 5. METHODS

### 5.1 Participants

28 participants took part in experiment I (18 – 49 years, mean age = 28.4, SD = 6.2; 9 female; 2 left-handed) and 20 participants took part in experiment II (21 – 49 years, mean age = 28.5, SD = 6.1; 9 female; 2 left-handed). All participants were fluent in English with self-reported normal hearing and participated voluntarily. The experiment was approved by the Regional Ethics Review Board in Stockholm (Dnr: 2017/998-31/2). Both written and oral informed consent were obtained from all participants prior to the experiments. All participants received a monetary compensation of SEK 600 (∼ EUR 60).

### 5.2 Quantification of individual musicality

A subset of the Goldsmiths musical sophistication index (MSI) self-report questionnaire^24,25^ containing 22 questions was used to estimate each participant’s level of musical sophistication. The MSI quantifies a participant’s musical skills, engagement, and behavior across multiple facets, making it ideal for testing a general population that includes both musicians and non-musicians. A copy of the questionnaire used for this study can be found in Supplementary Fig. S5. To ensure relevance to the selective auditory attention task, we emphasized on the Perceptual Ability, Musical Training, and Singing Abilities subscales. The Emotions subscale was excluded because our melody stimuli lack emotional content, rendering this subscale irrelevant to the task. Additionally, several questions from the Active Engagement subcategory were omitted to prioritize musical aptitude and ability over exposure. For instance, questions such as “Music is kind of an addiction for me – I couldn’t live without it,” which do not directly impact musical ability, were excluded. This specific combination of questions was previously used in a similar selective auditory attention study^23^ and the resultant MSI demonstrated strong correlations with performance. Among all participants, MSI scores ranged from 40 to 132, with a maximum possible score of 154 (m_I_ = 88.4, std_I_=26.8; m_II_ = 89.4, std_II_ = 28.9).

### 5.3 Task

For both experiments I and II, participants were presented with two simultaneous melodies of different pitch and instructed to selectively attend to either one, following a verbal cue. Each melody was composed of a series of 2 s tones and lasted for 8 - 26 s. During melody playback, participants were required to constantly follow the pitch contour of the attended melody until it stopped, at which point they reported the final direction of pitch change with a button press. The correct answer could be falling, rising or constant, corresponding to a chance level of 33 % (see Fig. 1). In each experiment, a total of 28 button responses were collected over approximately 10 minutes of MEG recording time for each participant. The fraction of correct button responses, out of 28, was computed as an index of task performance. To ensure that the sounds were separated based on features (in this case, pitch and timing) rather than location (i.e. left-versus-right), the stimuli were presented identically to both ears via insert earphones (model ADU1c, KAR Audio, Finland).

#### Experiment I

Tone onsets alternate between the low-pitched and high-pitched melodies, beginning with either melody (order balanced across trials). As each new tone would draw an effect of bottom-up attention towards it, this experimental design allowed us to dissociate top-down (from the cue) and bottom-up attention effects towards each melody.

#### Experiment II

Tone onsets for both melodies occurred concurrently, thereby engaging bottom-up attention simultaneously towards both melodies.

### 5.4 Stimuli

*Frequency-tagging*. The melodies were constructed with 2s long sinusoidal tones that were generated and amplitude-modulated using the Ableton Live 9 software (Berlin, Germany). The carrier frequencies of the low-pitched melody tones and high-pitched melody tones were 131 - 220 Hz and 329 – 523 Hz respectively. 256 tones were used to construct all 28 melodies of each pitch. For simplicity, only tones in the C major harmonic scale were used. Sinusoidal amplitude modulation of the tone was carried out by modifying its amplitude envelope, which corresponds to its raw sound volume, with regular increases and decreases according to a sine wave (see Supplementary Fig. S7). The rate of modulation, or modulation frequency (f_m_), was 39 Hz for the low-pitched melody and 43 Hz for the high-pitched melody. The modulation frequency was centered around 40 Hz and the modulation depth (m) was set at 100% to maximize cortical ASSR power^1^.

#### Stimulus presentation

The melody duration was randomized to last for 8 - 26 s each, preventing participants from predicting when it ends and thus maintaining their high levels of attention throughout playback. The volume of the high-pitched melody was reduced by 10 Db relative to the low-pitched melody to balance subjective loudness differences across frequency ranges^63^. Identical stimuli were delivered to both ears through sound tubes, with the volume calibrated to approximately 75 dB SPL per ear using a sound meter (Type 2235, Brüel & Kjær, Nærum, Denmark) and slightly adjusted for individual comfort. To ensure that the stimulus was presented with less than 1 ms jitter, we used a specialized auditory setup comprising of the AudioFile Stimulus Processor (Cambridge Research Systems Limited) coupled with customized scripts in Presentation® (Version 18.3, Neurobehavioral Systems, Inc., Berkeley, CA, www.neurobs.com).

### 5.5 Magnetoencephalography data acquisition & preprocessing

MEG measurements were carried out using a 306-channel whole-scalp neuromagnetometer system (Elekta TRIUXTM, Elekta Neuromag Oy, Helsinki, Finland). Data was recorded at a sampling rate of 1 kHz, bandpass filtered onilne between 0.1 - 330 Hz, and stored for offline analysis. Horizontal eye-movements and eye-blinks were monitored using horizontal and vertical bipolar electrooculography electrodes. Cardiac activity was monitored with bipolar electrocardiography electrodes attached below the left and right clavicle. Internal active shielding was active during MEG recordings to suppress electromagnetic artefacts from the surrounding environment. In preparation for the MEG-measurement, each participant’s head shape was digitized using a Polhemus Fastrak. The participant’s head position and head movement were monitored during MEG recordings using head-position indicator coils. The acquired MEG data was pre-processed using MaxFilter^64,65^ by applying temporal signal space separation (tSSS) to suppress artefacts from outside the MEG helmet and compensate for head movement during the recording, before being transformed to a default head position. The tSSS had a buffer length of 10 s and a cut-off correlation coefficient of 0.98.

### 5.6 MEG data analysis

Following preprocessing, subsequent MEG data analyses were done using the MNE-Python software package^66^. Firstly, we applied a 1 - 50 Hz bandpass filter to the continuous MEG data. Next, eye movements and heartbeats artefacts were automatically detected from electrooculographic and electrocardiography signals using independent components analysis (ICA, fastICA algorithm), validated visually, and removed for each participant. On average, we removed 4.9 components (SEM 0.2) across participants.

#### Experiment I data analysis

The continuous MEG data was segmented into 1 s long epochs from each tone onset. Since the salient tone onsets alternate between the low- and high-pitched melodies, bottom-up attention was strongly drawn towards either melody tone but never both. Thus, we defined all epochs into two conditions based on whether bottom-up attention was directed to the low- or high-pitched melody tone. Alternatively, we also defined all epoches into another two conditions based on whether the cue directed top-down attention to the low- or high-pitched melody. In both instances, 228 epochs were defined per condition per participant.

#### Experiment II data analysis

The first 0 - 1 s from each tone onset was defined as the early half of the tone, while the subsequent 1 - 2 s was defined as the late half of the tone, defining two conditions by time. Subsequently, each of the two time conditions were analyzed separately for top-down attention according to the cue, categorizing the epochs into attend low- or high-pitched for each of the early and late half. As a result, about 114 epochs were defined per condition per participant.

### 5.7 Classification of attention

*Repeated-splitting support vector machine*. To classify attention conditions at single-subject level, we designed a specialized “repeated-splitting” decoder that was highly sensitive to ASSRs. These customized algorithms were based on the scikit-learn package in python. For each condition, epochs were randomly divided into five groups and averaged within each group to produce five evoked ASSRs. Next, the five evoked ASSRs were fast fourier transformed (frequency resolution = 1 Hz) to acquire five power spectra. Spectral smoothing was performed using a Hann window in experiment I and a boxcar window in experiment II. For any contrast between two conditions, the corresponding power spectra (5 each condition) were fed into a binary linear support vector machine with all 306 sensors as features, and classified using a 5-fold cross validation method. We computed the area under the receiver operating characteristic curve (AUC) as a measure of discriminability between conditions (chance level = 0.5). Finally, we repeated the entire process from the initial group division step 1000 times and computed the mean AUC across repetitions. The AUC was computed independently for each frequency from 4 - 45 Hz. Fig. 2a outlines each of the steps described above.

For *experiment I*, the AUC across frequency was then averaged across 25 participants (1 participant was excluded due to incomplete data collection, and another 2 participants were excluded due to excessive noise at sensor-level; see Supplementary Fig. S8) to give the group-averaged AUC across frequency. For each participant in *experiment II*, the AUC across frequency was computed separately for the early and late half of the tone, before combining them by selecting the higher AUC between the two halves at each frequency. The resultant “best of two” AUC across frequency was then averaged across all 20 participants to give the group-averaged AUC across frequency.

#### Sliding window analysis

To investigate fluctuations in selective attention across time during experiment II, we employed a sliding window approach (fast fourier transform, window length = 1 s, step size = 50 ms, boxcar window) across the 2 s tone duration for the ASSRs, computing the power specifically at 39 and 43 Hz. Classification was performed independently for each time window, resulting in the AUC across time at 39 and 43 Hz, corresponding to the attentional modulation of the low- and high-pitched melody respectively. For each participant, we determined the time of peak selective attention during which the AUC was maximum, regardless of which melody it corresponded to. Participants were split into two equal groups of ten based on their time of peak selective attention, which was before 0.5 s for early attendees and after 0.5 s for late attendees. Subsequently, we compared differences in musicality and performance between the early and late attendees using a two-tailed permutation test with 10 000 shuffles.

### 5.8 Structural Magnetic Resonance Imaging acquisition and processing

Anatomical MRIs were acquired using hi-res Sagittal T1 weighted 3D IR-SPGR (inversion recovery spoiled gradient echo) images by a GE MR750 3 Tesla scanner with the following pulse sequence parameters: 1 mm isotropic resolution, FoV 240 × 240 mm, acquisition matrix: 240 × 240, 180 slices 1 mm thick, bandwidth per pixel=347 Hz/pixel, Flip Angle=12 degrees, TI=400 ms, TE=2.4 ms, TR=5.5 ms resulting in a TR per slice of 1390 ms. Cortical reconstruction and volumetric segmentation of all participants’ MRI was performed with the Freesurfer image analysis suite^67^.

### 5.9 MEG source reconstruction

Source localization of the MEG signals (combined planar gradiometers and magnetometers) was performed using a minimum-norm estimate^68^ (MNE) distributed source model containing 20484 dipolar cortical sources. The MEG signals were coregistered onto each individual participant’s anatomical MRIs using previously digitized points on the head surface during MEG acquisition. Based on our previous studies^23,26,27^ on ASSR sources, we demarcated six regions of interest, namely the orbital gyrus (OrG), superior temporal gyrus (STG), and inferior parietal lobe (IPL) at each hemisphere, according to the Brainnetome Atlas^28^.

### 5.10 Classification of selective attention (source level)

To classify attention conditions at individual source level, we incorporated the source localization step into the “repeated-splitting” support vector machine decoder (with identical parameters previously described in 7 Classification of attention) as follows: For each contrast, epochs per condition were randomly divided into five groups and averaged within each group to produce five evoked ASSRs. The five evoked ASSRs were then source localized to produce five source estimations across the entire cortex. For each of the six regions of interest (ROIs), a subset of these sources was selected to contain only sources originating within the ROI. The data was then fast fourier transformed. The power at 39 and 43 Hz was combined in feature space before being classified into one of two conditions as we measured the decoding performance with the AUC. Finally, we repeated the entire process from the initial group division step 1000 times and computed the mean AUC across repetitions. One participant was excluded due to incomplete data collection in both experiments, resulting in a final sample of 27 participants in experiment I and 19 participants in experiment II for all source-level analysis.

#### Lateness Index

For experiment II, we compared the AUC between the early and late tone half to investigate fluctuations in selective attention across time at source level. This was done in place of the sliding window approach at sensor level to reduce computational costs. For each participant, we computed a lateness index using the formula: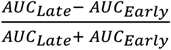 to quantify the relative strength of selective attention in the late compared to the early tone half. Following which, we carried out Pearson correlation tests between the lateness index and MSI, as well as between the lateness index and task performance to examine which brain regions exhibit relationships between lateness and these behavioral measures. We reported two-tailed p values from these tests. Data normality was assessed using the Shapiro-Wilk test at a threshold of p < 0.05 for non-normal distributions and visually confirmed through quantile–quantile plots.

## 7. ACKNOWLEDGEMENTS

Data for this study was collected at the Swedish National Facility for Magnetoencephalography, Karolinska Institutet, Sweden. This facility is supported by Knut and Alice Wallenberg Foundation (KAW2011.0207). The present studies were supported by the Swedish Foundation for Strategic Research (SBE 13-0115). The computations for source analysis were enabled by the Berzelius resource provided by the Knut and Alice Wallenberg Foundation at the National Supercomputer Centre.

## 8. AUTHOR CONTRIBUTIONS

C.M. and D.L. designed the study, set up the experiments and wrote the manuscript. C.M. collected and analyzed the data, with valuable input from D.L., D.P., and J.G.

## 9. COMPETING INTERESTS

The authors declare no competing interests.

## 10. DATA AND MATERIAL AVAILABILITY

All data needed to evaluate the conclusions in the paper are present in the paper and/or the Supplementary Materials. All code is publicly available at https://github.com/cassialmt/ALTOL_ms

## Notes

### Competing Interest Statement

The authors have declared no competing interest.

### Summary of Updates

Abstract revised; Figures 1 - 5 revised, adding mini brains inline to visualize regions-of-interest; Introduction revised to add content on previous ASSR work; Discussion revised;

