## supplementary material for "How musicality enhances top-down and bottom-up selective attention: Insights from precise separation of simultaneous neural responses"

Science Advances
AAAS

Supplementary Materials for

### How musicality enhances top-down and bottom-up selective attention: Insights from precise separation of simultaneous neural responses

Cassia Low Manting* *et al.*

**This PDF file includes:**

Figs. S1 to S8
Tables S6

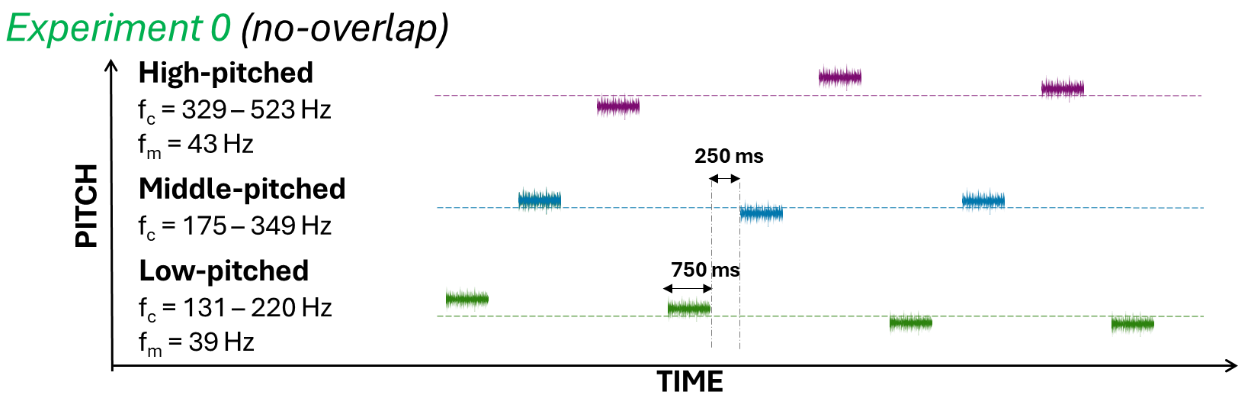

**Fig. S1 | Experiment 0 design: No overlap.** Participants were listened to three melody streams that differed in pitch, attending to either the low- or high-pitched melody and reporting the direction of pitch change at the end. Throughout stimulus playback, the tone onsets followed a consistent pattern alternating from low- to middle- to high-pitched (as in the above example) or its reverse. The modulation frequencies (fm) and carrier frequency (fc) ranges of the low- and high-pitched melodies were identical to experiments I and II. In addition to the middle-pitched melody, which was used as a reference, each tone duration is 750 ms instead of 2 s, with the introduction of 250 ms of silence between tones. This experiment is considered the least complex as the melodies are most easily distinguishable without any overlap in time.

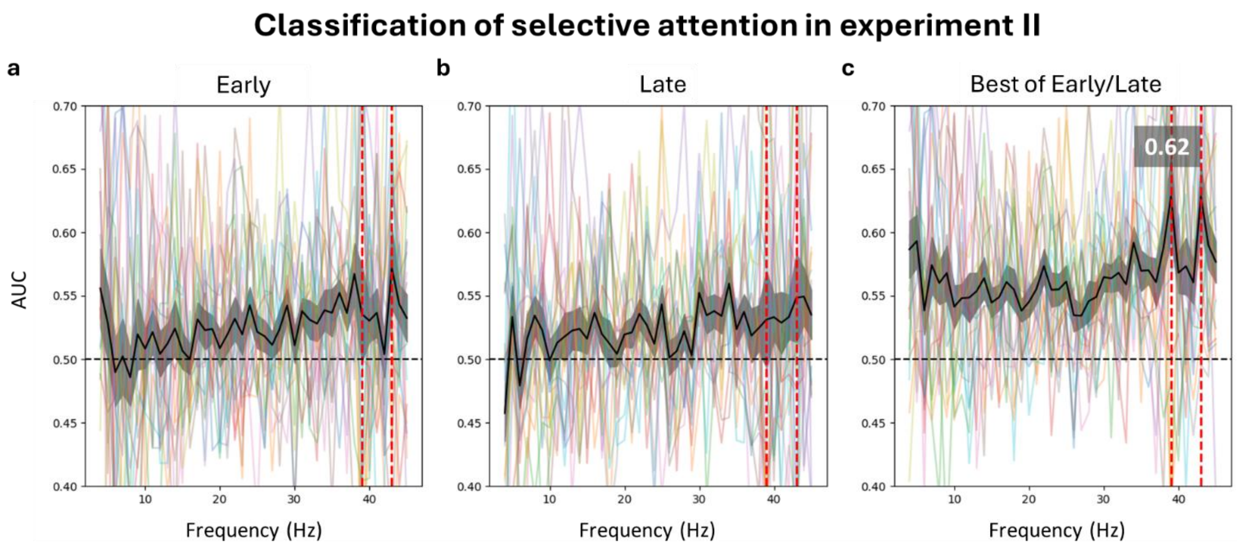

**Fig. S2 | Classification of selective attention in experiment II.**
**a** and **b**, Across participants (N = 20), the classification between attention conditions did not show any peaks at the expected modulation frequencies (i.e. 39 and 43 Hz) in the early or late half of the 2s tone. **c**, 39 Hz and 43 Hz discrimination peaks (AUC = 0.62 for 39 and 43 Hz) were observed when taking the higher score between the two halves at each frequency. This indicates that although selective attention was active across participants, there were variations in when and how it occurred over time. Red vertical lines mark the AUC at 39 and 43 Hz. Shading indicates s.e.m.

### Experiment I: Correlations of selective attention with musicality & performance

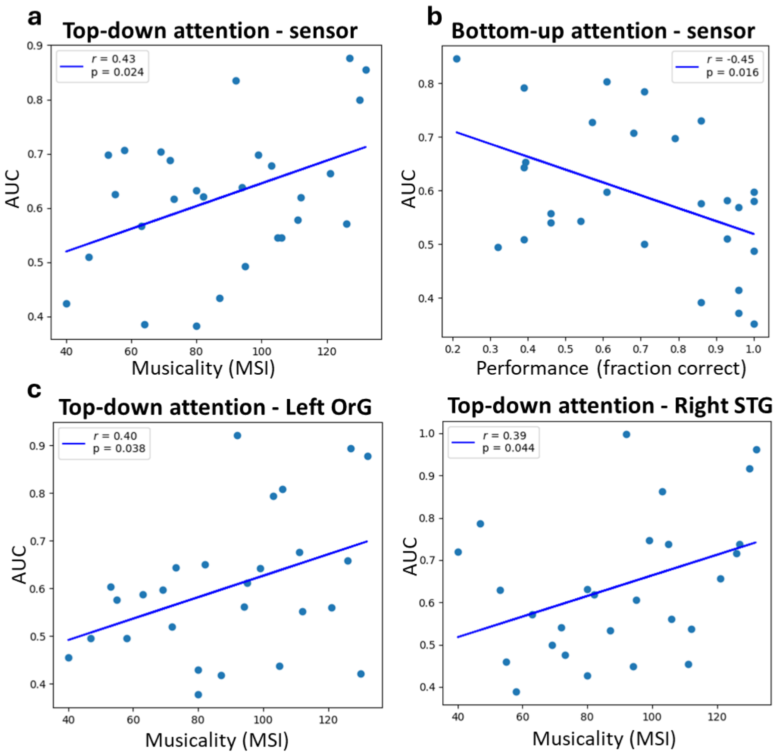

**Fig. S3 | Correlations of selective attention with musicality & task performance in experiment I.**
Scatterplots showing correlation between classifier area-under-curve (AUC) values against individual musicality or task performance. The AUC (y-axis) reflects the degree of selective attention. At sensor level, top-down attention correlates positively with musicality (N = 28, Pearson correlation, r = 0.42, $p < 0.05$) in **a,** while bottom-up attention correlates negatively with performance (N = 28, Pearson correlation, r = 0.45, $p < 0.05$) in **b**. For **a** and **b**, the y-axis plots the mean AUC between 39 and 43 Hz. **c**, At source level, top-down attention correlated positively with musicality at the left OrG (N = 27, Pearson correlation, r = 0.40, $p < 0.05$) and right STG (N = 27, Pearson correlation, r = 0.39, $p < 0.05$).

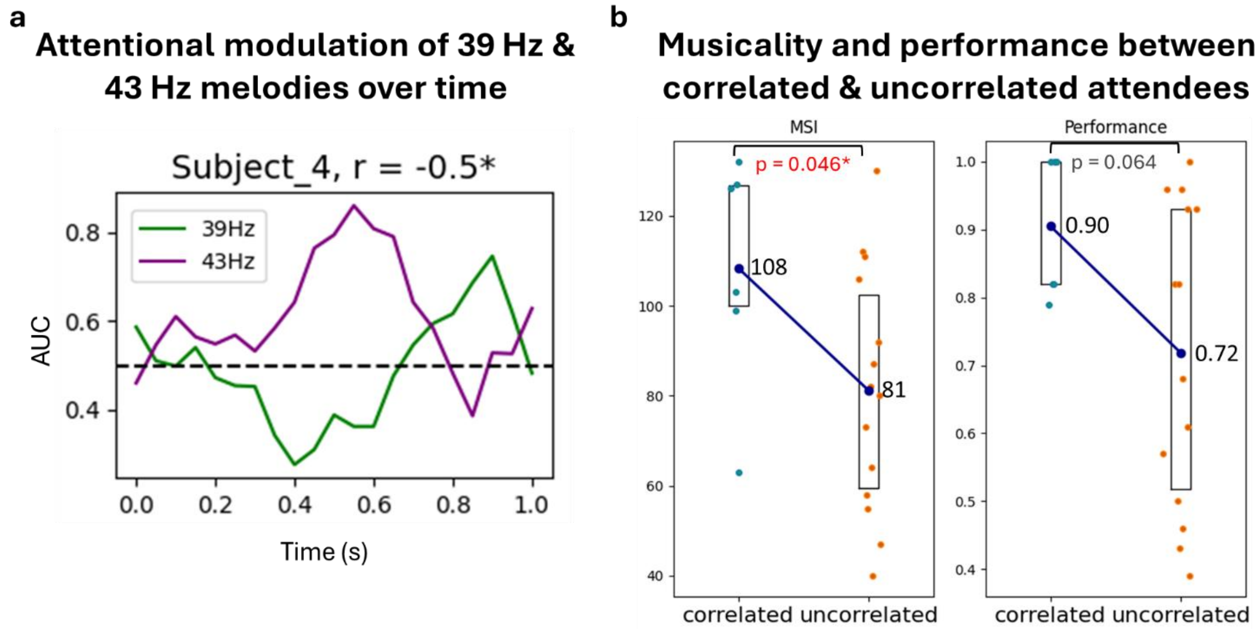

**Fig. S4 | Tracking attentional modulation of each melody across tone duration.**
**a**, We employed a 1s sliding time window to track how attentional modulation of the 39 (green) and 43 (purple) Hz frequency-tagged melodies each vary across the 2s tone duration. **b,** In six participants (correlated attendees), significant correlations were found between the attentional modulation of the two melodies across time. These participants were more musical ($p < 0.05$, two-tailed permutation test) than the remaining 14 participants (uncorrelated attendees), indicating that musical participants might have employed organized strategies to manage attention towards simultaneous sounds. In the above example, the correlation was negative, suggesting that the participant employed a strategy that involved switching attention between melodies. Box plots represent the 25th – 75th percentiles of the data, with the center dot indicating the mean.

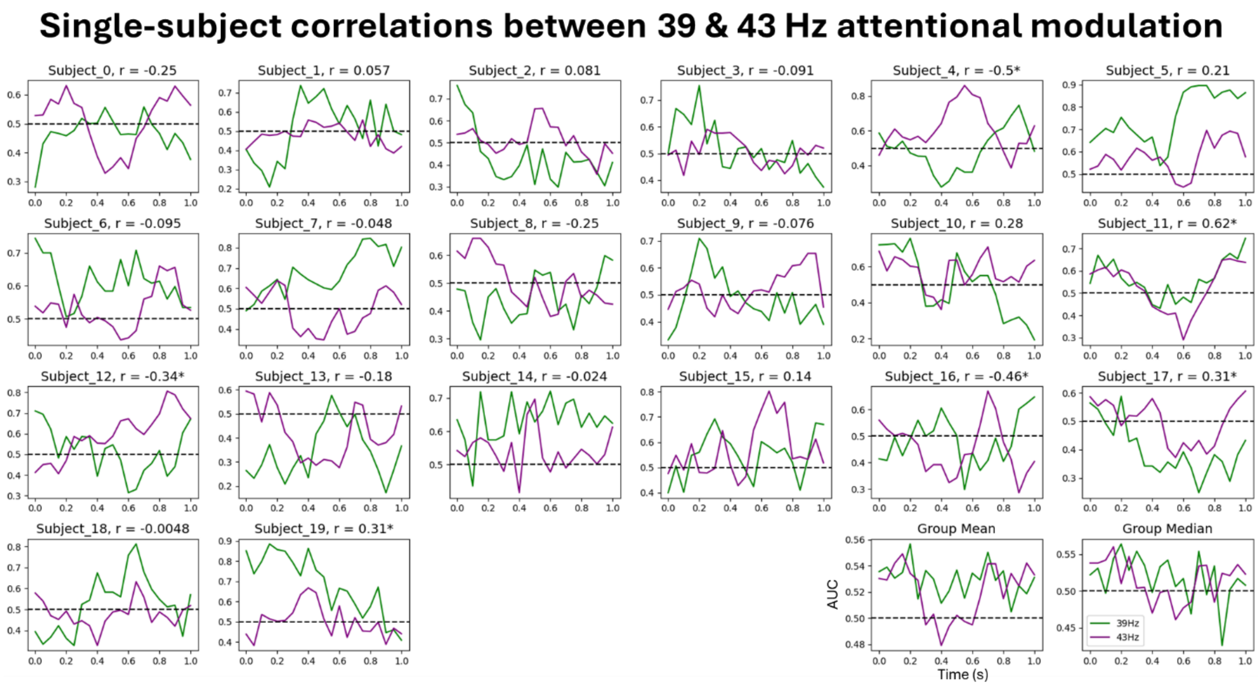

**Fig. S5 | Single-subject correlations between attentional modulation of each melody.** Significant correlations (Kendall Tau, $p < 0.05$, marked by asterisks) between the 39 Hz (green) and 43 Hz (purple) attentional modulation curves only emerged in six subjects. For several other participants (e.g. Subject_5), the curves exhibit more complex correlations, including time lags, which cannot be effectively characterized by the Kendall's Tau. For all subplots, the x-axis and y-axis denote time and AUC respectively. Dashed lines indicate 0.5 chance level. The last two subplots in the bottom-right corner show the mean and median across all 20 subjects for each curve.

**Table S6 | Subset of Goldsmiths Musical Sophistication Index questionnaire.**

| |
|---|
| ***Please select from:*** |
| ***1 Completely Disagree/ 2 Strongly Disagree/ 3 Disagree/ 4 Neither Agree nor Disagree/ 5 Agree/ 6 Strongly Agree/ 7 Completely Agree*** |
| 1. I spend a lot of my free time doing music-related activities. |
| 2. If somebody starts singing a song I don't know, I can usually join in. |
| 3. I am able to judge whether someone is a good singer or not. |
| 4. I usually know when I'm hearing a song for the first time. |
| 5. I can sing or play music from memory. |
| 6. I am able to hit the right notes when I sing along with a recording. |
| 7. I find it difficult to spot mistakes in a performance of a song even if I know the tune. |
| 8. I have trouble recognizing a familiar song when played in a different way or by a different performer. |
| 9. I have never been complimented for my talents as a musical performer. |
| 10. I am not able to sing in harmony when somebody is singing a familiar tune. |
| 11. I can tell when people sing or play out of time with the beat. |
| 12. I can tell when people sing or play out of tune. |
| 13. When I sing, I have no idea whether I'm in tune or not. |
| 14. I would not consider myself a musician. |
| 15. After hearing a new song two or three times, I can usually sing it by myself. |
| 16. I only need to hear a new tune once and I can sing it back hours later. |
| *Please select from the underlined options:* |
| 17. I engaged in regular, daily practice of a musical instrument (including voice) for _0/1/2/3/4-5/6-9/10 or more_ years. |
| 18. At the peak of my interest, I practiced _0/0.5/1/1.5/2/3-4/5 or more_ hours per day on my primary instrument. |
| 19. I have had formal training in music theory for _0/0.5/1/2/3/4-6/7 or more_ years |
| 20. I have had _0/0.5/1/2/3-5/6-9/10 or more_ years of formal training on a musical instrument (including voice) during my lifetime. |
| 21. I can play _0/1/2/3/4/5/6 or more_ musical instruments |
| 22. I listen attentively to music for _0-15 min/ 15-30 min/ 30-60 min/ 60-90 min/ 2 h/ 2-3 h/ 4 h or more_ per day. |
| *Additional question not used for computing MSI:* |
| 23. The instrument I play best (including voice) is ____ |

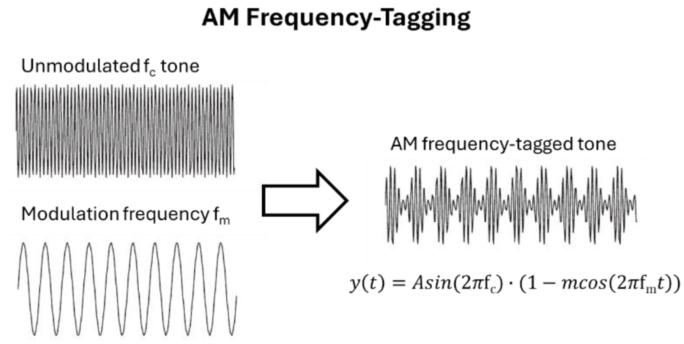

**Fig. S7 | Sinusoidal amplitude modulation (AM) frequency-tagging of a tone.** The amplitude envelope of the original unmodulated tone is modulated systematically according to the modulation frequency ($f_m$). This is equivalent to increasing and decreasing the sound volume at a rate equal to $f_m$. The degree of modulation is adjusted by the modulation depth (m). The resultant frequency-tagged tone is described by the displayed equation where A and $f_c$ corresponds to the amplitude and carrier frequency of the unmodulated tone. The above image was edited from Luo 2006[1].

1. Luo, H., Wang, Y., Poeppel, D. & Simon, J. Z. Concurrent Encoding of Frequency and Amplitude Modulation in Human Auditory Cortex: MEG Evidence. *J. Neurophysiol.* **96**, 2712–2723 (2006).

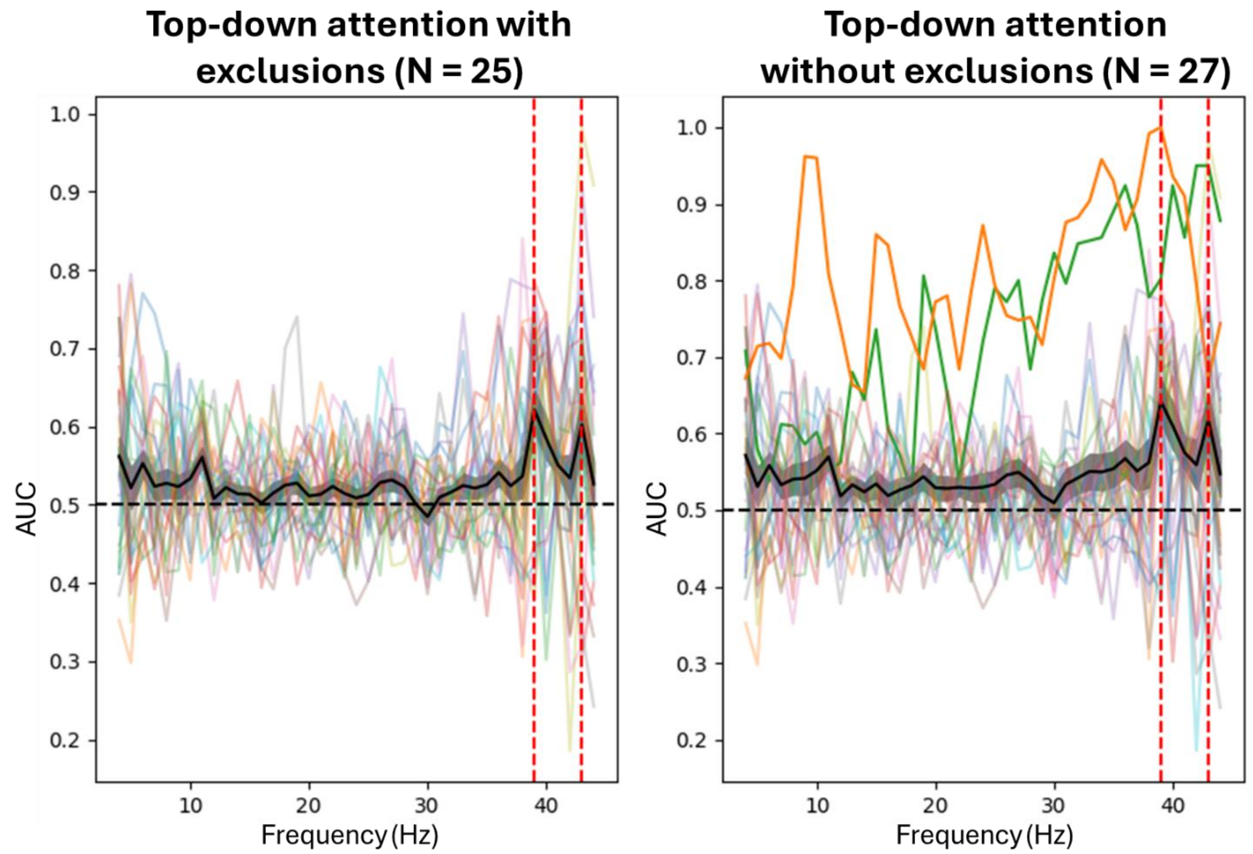

**Fig. S8 | Butterfly plots across participants for top-down attention in experiment I with (left) and without (right) participant exclusions due to excessive noise**. The mean across participants is depicted in black, with red vertical lines marking the AUC peaks at 39 and 43 Hz. Shading indicates s.e.m. In the right figure, excluded participants are shown with full opacity (orange and green) to emphasize their higher noise levels compared to the remaining participants (30% opacity).
